# Geographical gradients in leaky sex expression and reproductive effort in a dioecious plant are consistent with selection during range expansion

**DOI:** 10.64898/2026.01.13.699234

**Authors:** M. T. Nguyen, J. R. Pannell

## Abstract

Range expansion involves repeated dispersal and the establishment of new populations through colonization. Natural selection is therefore expected to favour traits that enhance dispersal and enable uniparental reproduction when mates are scarce. It is known that range-edge populations may evolve increased dispersal and, in hermaphroditic species, an enhanced capacity for self-fertilization. Here, we report results from a large common-garden experiment showing that males and females of the dioecious annual herb *Mercurialis annua* are more likely to show ‘leaky’ or inconstant sex expression towards the recently colonized edge of its geographical distribution in Europe than those from the eastern Mediterranean Basin, from where the species expanded its range. Range-edge populations also had greater reproductive effort (RE). Leaky sex expression allows otherwise unisexual individuals of *M. annua* to produce progeny by self-fertilization, a trait that may have been selected for reproductive assurance during the range expansion. High RE likely facilitated both colonization and subsequent population establishment. Our results show that selection during range expansions may modify reproductive traits towards range margins, extending this notion to the phenomenon of leaky sex expression in dioecious species. Previous work has shown that leaky sex expression in *M. annua* is strongly heritable; our results now further suggest that has been enhanced by selection for reproductive assurance towards range margins and confirm, in a natural setting, that the trait is not just an expression of developmental instability during floral development or a relic of an incomplete transition to dioecy.

## Introduction

The current distributions of many species, especially at mid to high latitudes, are the legacy of past range expansions, often from Pleistocene refugia (Hewitt, 2000; Svenning & Skov, 2007). Such expansions leave characteristic population-genetic signatures, both because populations encounter new abiotic and biotic environments and because repeated colonization entails serial bottlenecks (Bridle & Vines, 2007; Sexton *et al*., 2009). These bottlenecks increase drift and progressively erode genetic diversity with distance from the ancestral refugium, affecting both neutral and fitness-related variation (Ibrahim *et al*., 1996; Hewitt, 2000; Excoffier *et al*., 2009). As a result, range expansion can lead to the accumulation of deleterious mutations, changes in homozygosity and inbreeding depression, and a reduced efficacy of selection towards the range edge (Pujol *et al*., 2009; Peischl *et al*., 2013, 2015; González-Martínez *et al*., 2017; Willi *et al*., 2018). At the same time, colonization itself should favour phenotypes that enhance establishment and spread, so that populations towards a species’ range margin may evolve traits distinct from those in the core range and former refugia. The evolution of such traits is likely to be an important component of adaptation at range margins.

A key idea is that individuals capable of uniparental reproduction, whether by asexual means (Delnevo *et al*., 2024) or self-fertilization (Pannell, 2015), should be more likely to succeed in colonizing new habitats than those that need to find a mate to reproduce. Baker (1955) linked observations of an increased capacity for self-fertilization in species on oceanic islands to colonization following long-distance dispersal to islands. This enrichment on islands of species with a capacity to self-fertilize as a result of selection for reproductive assurance was later labelled Baker’s Law by Stebbins (1957), highlighting its broad relevance across both plants (Baker, 1955, 1967) and animals (Longhurst, 1955; Stebbins, 1957). Substantial further evidence for Baker’s Law has accumulated since Baker’s initial paper (Grossenbacher *et al*., 2017), and the concept has been expanded to explain the evolution of a capacity for uniparental reproduction in species with a colonizing habit in metapopulations with frequent population turnover (Pannell, 1997, 2000; Pannell & Barrett, 1998) as well as in the context of species range expansions (reviewed in Cheptou, 2012; Pannell, 2015; Pannell *et al*., 2015). In metapopulations, higher extinction rates and greater patchiness are associated with increased frequencies of self-fertilizing individuals in species with mixed mating, including *Silene vulgaris* (Taylor *et al*., 1999), *Helicteres brevispira* (Franceschinelli & Bawa, 2000), *Eichhornia paniculata* (Barrett & Husband, 1990; Ness *et al*., 2010) and *Mercurialis annua* (Obbard *et al*., 2006b). In the context of range expansion, autonomously apomictic species tend to occupy broader distributions than their sexually reproducing relatives (Hörandl *et al*., 2008; Cosendai *et al*., 2013; Hörandl, 2023), and marginal populations are more likely to possess the ability to reproduce by self-fertilization (Lloyd, 1965; Moeller & Geber, 2005; Obbard *et al*., 2006b; Busch & Delph, 2012; Ruane *et al*., 2020). For example, range-margin populations of *Clarkia xantiana* ssp. *parviflora* show significantly lower herkogamy, petal area, and protandry than core populations, indicating increased selfing ability, a pattern further supported by emasculation experiments (Ruane *et al*., 2020). Similarly, in *Leavenworthia* (Cruciferae), floral morphology and extensive field surveys suggest a breakdown of self-incompatibility in range-marginal populations (Lloyd, 1965).

Selection on traits that affect colonization ability should also apply to dioecious species, as originally discussed by Baker (1967). In the context of Baker’s Law, the frequently observed enrichment of dioecious species on islands (Carlquist, 1965, 1967) might appear surprising, given that dioecy implicates biparental rather than uniparental reproduction. However, Baker (1967) already drew attention to the fact that the males and females or many dioecious plant species produce occasional flowers with the opposite sexual function, potentially allowing them to self-fertilize. This phenomenon, referred to as inconstancy or ‘leakiness’ in sex expression (e.g., Baker, 1967; Lloyd & Bawa, 1984; Korpelainen, 1998), is common in dioecious species (Ehlers & Bataillon, 2007; Cossard & Pannell, 2019). Leakiness in sex expression has tended to be attributed to developmental noise (Lloyd & Bawa, 1984), a plastic response to environmental cues such as temperature, drought, herbivory (Charnov & Bull, 1977; Freeman *et al*., 1980; Case & Barrett, 2001, 2004; Delph, 2003; Delph & Wolf, 2005; Golenberg & West, 2013; Villamil *et al*., 2022), or as evidence of still incomplete transitions from combined to separate sexes (Charlesworth & Charlesworth, 1978; Lloyd, 1980; Ross, 1980, 1982). Ehlers and Bataillon (2007) and Crossman and Charlesworth (2014) modelled the evolution and maintenance of leakiness in terms of a response to selection for reproductive assurance, and Cossard et al. (2021) and Gerchen et al. (Gerchen *et al*., 2026) have recently demonstrated that variation in leakiness can have a genetic basis and be responsive to selection. However, empirical evidence that leakiness can be promoted specifically by selection for reproductive assurance such as in metapopulations or range expansions is still lacking.

Selection during range expansion might also bring about evolution of dispersal and life-history traits that promote both colonization and rapid establishment (reviewed in Hargreaves & Eckert, 2014; Chuang & Peterson, 2016; Weiss-Lehman & Shaw, 2022). Increased dispersal may be selected directly through the process of colonization itself (Travis & Dytham, 1999) and may be further reinforced by assortative mating among highly dispersive individuals at the expansion front (Shine *et al*., 2011). A notable example is provided by the invasive cane toad *Rhinella marina* in Australia, which has evolved longer legs enabling faster dispersal at the invasion front (Phillips *et al*., 2006, 2007). In plants, dispersal-related traits such as enhanced seed pappus or wing development have been observed in recently colonised compared to older populations (Cody & Overton, 1996; Cheptou *et al*., 2008; Darling *et al*., 2008; Monty & Mahy, 2010), a phenomenon that has been labelled the ‘metapopulation effect’ (Olivieri & Gouyon, 1997) but that also applies to range expansions. For instance, the ragwort *Senecio inaequidens* exhibits increased pappus size, enhancing wind-dispersal distance toward its range margin (Monty & Mahy, 2010), and the dune plant *Abronia umbellata* shows greater seed wing index in more marginal populations, a legacy of selection on colonization ability (Darling *et al*., 2008). Successful population establishment then should be enhanced greater allocation to reproduction, at the expense of traits associated with persistence and competitive ability in crowded environments, such as vegetative growth, size and competitive performance (Tilman, 1990; Burton *et al*., 2010; Urquhart-Cronish *et al*., 2024). The expected shifts along this trade-off with range expansion accord, more generally, with classic life-history ideas, including the colonization-competition trade-off (Tilman, 1990), the r/K continuum (MacArthur & Wilson, 1967; Pianka, 1970; Reznick *et al*., 2002; Travis *et al*., 2023), and Grime’s C–S–R framework (Grime, 1974), which predict that populations experiencing repeated colonization should evolve traits associated with more ruderal, colonizing strategies (MacArthur & Wilson, 1967; Pianka, 1970; Grime, 1974; Stearns, 1983, 1989; Reznick *et al*., 2002; Travis *et al*., 2023). Reproductive effort, the proportion of biomass allocated to flowering and fruiting, is an easily measurable trait corresponding to this trade-off that should accordingly increase with range expansion.

Here, we test the hypothesis that selection should have favoured increased leakiness in sex expression and greater reproductive effort in populations towards the margins of the geographic range of the diploid European annual dioecious plant, *Mercurialis annua* L. (Euphorbiaceae). *M. annua* is a wind-pollinated annual colonizer of disturbed habitat that expresses leakiness in sex expression in natural populations for both males and females. The rapid evolution of enhanced leakiness in experimental populations (Cossard *et al*., 2021; Gerchen *et al*., 2026) indicates that the trait is strongly heritable. Our study involved sampling dioecious *M. annua* from populations across its geographic range in Europe, growing them in a common garden, and measuring the size of plants and their allocation to male and female flowers and fruits. We used our measures to quantify the leakiness in sex expression and the reproductive effort of all individuals sampled. Leakiness can be measured both by its incidence (the proportion of individuals producing opposite-sex flowers) and its degree (the number of opposite-sex flowers produced per individual) (Cossard & Pannell, 2019). We expected reproductive assurance to favour increased incidence of leakiness in both sexes with distance from the ‘expansion origin’ (see Methods for its definition), but an increased degree of leakiness only in males. This is because a mildly leaky female could self-fertilize many ovules and establish a population, whereas the success of a colonizing male would depend on it producing many seeds, requiring a substantial degree of leakiness. An greater degree of leakiness in males than females is also expected as an outcome of selection towards female-biased sex allocation due to local mate competition (Hamilton, 1967; Roux *et al*., 2024).

## Methods

### Study species and population sampling

*Mercurialis annua* L. (Euphorbiaceae) is a European, dioecious, wind-pollinated annual plant with a distribution from the eastern Mediterranean through central and western Europe to the Atlantic Ocean. The species occupies disturbed ruderal sites and has a life-history typical of weedy colonizing annuals and a demography characterised by metapopulation dynamics (Obbard *et al*., 2006b; Eppley & Pannell, 2007b; Pannell *et al*., 2008; Pujol *et al*., 2009; Pannell *et al*., 2014; Dorken *et al*., 2017; González-Martínez *et al*., 2017). Populations of *M. annua* in western Europe have lower within-population genetic diversity than those in the east (Obbard *et al*., 2006b) and show signs of increased genetic load and with fewer genomic signatures of selective sweeps than those in the east (González-Martínez *et al*., 2017), consistent with a range expansion into western Europe from a Pleistocene refugium in the eastern Mediterranean Basin.

*M. annua* possesses heteromorphic sex chromosomes with an XY sex-determination system dated back to over 1 MY ago (Russell & Pannell, 2015; Veltsos *et al*., 2018, 2019; Gerchen *et al*., 2022) and strong sexual dimorphism (Harris & Pannell, 2008; Hesse & Pannell, 2011; Sánchez Vilas & Pannell, 2011; Tonnabel *et al*., 2019). It also shows substantial leakiness in sex expression in both sexes (Cossard & Pannell, 2019; Cossard *et al*., 2021). Flowering in *M. annua* is indeterminate. Females and males of *M. annua* produce pistillate flowers and staminate inflorescences, respectively, in the axis of each new leaf. Leaky females usually produce one or more staminate flowers alongside its axillary female flowers. Leaky males produce pistillate flowers and fruits along with their staminate flowers on extended inflorescence stalks (‘peduncles’). Both males and females with leaky sex expression are capable of self-fertilizing (Eppley & Pannell, 2007a; Li *et al*., 2019; Cossard *et al*., 2021) and thus enjoy reproductive assurance in the absence of mates of the opposite sex, e.g., during colonization or in sparse stands (Baker, 1967; Pannell *et al*., 2008, 2015; Cheptou, 2012; Cossard *et al*., 2021).

For our study, we collected seeds, or had them sent to us, from 26 populations across the distribution of diploid *M. annua* in autumn 2023 and spring 2024 (Figure 1, Table S1). For each population, seeds were bulked from at least 20 females, except for the Albanian population, where only five individuals could be sampled due to its small size. In autumn 2024, we sowed seeds from all 26 populations in a peat-based substrate in trays set up on a glasshouse at the University of Lausanne, Switzerland. After one month, when most plants had begun flowering, we transplanted between 80 and 120 individuals per population into 0.68 L pots filled with Ricoter 140 soil (2,694 plants in total). Exceptions were Populations 1 and 8, where only 43 and 55 plants germinated, respectively, and all of these were used.

**Figure 1.**
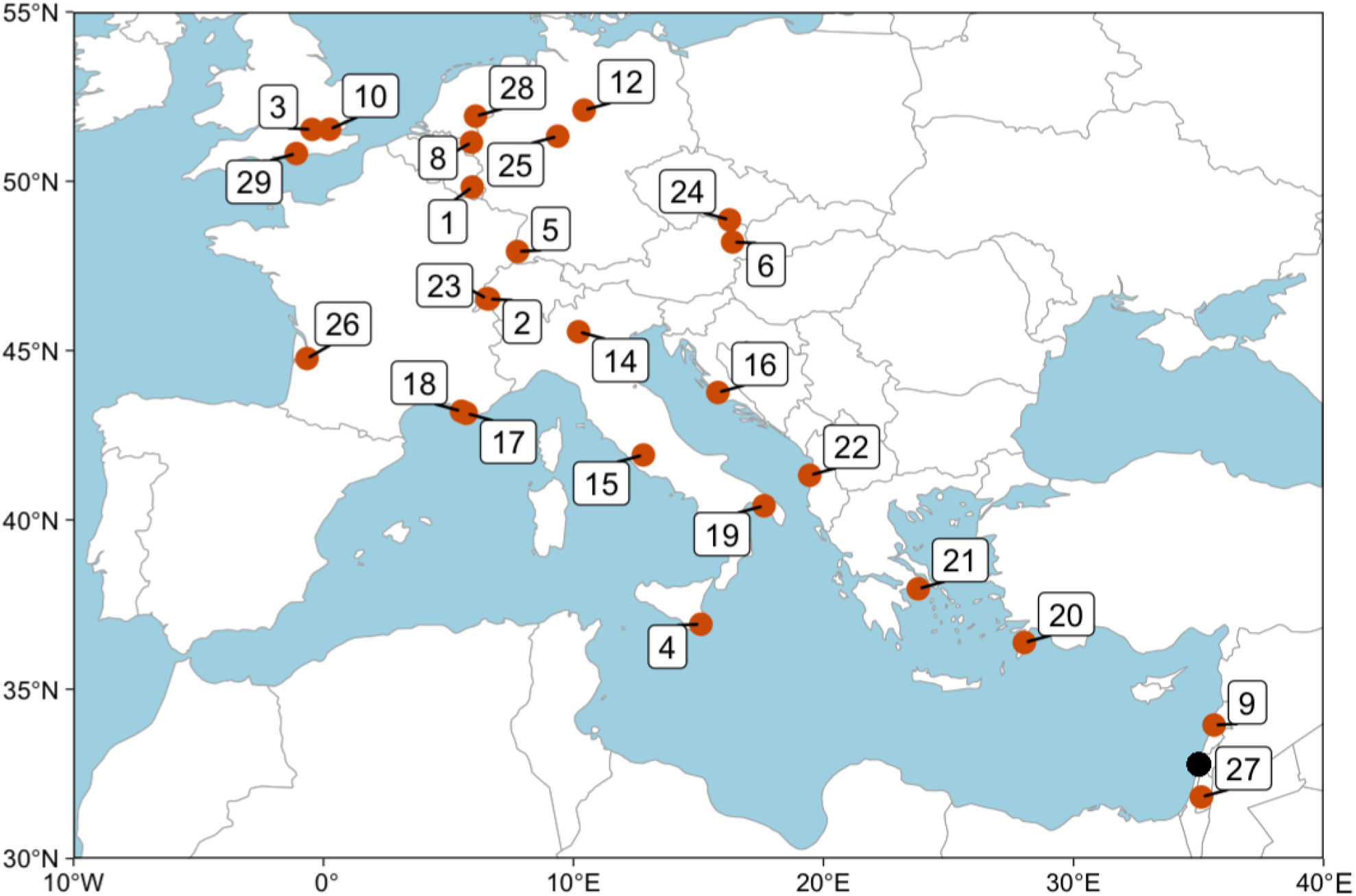
Map of sampling sites in Europe (red points) with their labels (detailed in Table S1) The point of origin (black point) assumed in our analyses for the expansion origin is midway between populations 9 and 27.

### Experimental treatments and design of the common garden

Sex expression in *M. annua* is mediated by the plant hormones auxin (masculinizing) and cytokinin (feminizing) (Louis & Durand, 1978; Dauphin-Guerin *et al*., 1980; Hamdi *et al*., 1987; Li *et al*., 2019), and exogenous application of these hormones can bring about altered sex expression, with cytokinin application causing males to produce female flowers and fruits, and auxin causing females to produce male flowers. Because we were concerned that we might have limited power to test our main hypothesis about leakiness if too few plants expressed it, to amplify levels of leakiness we applied hormone treatments to half of the plants in 24 of the populations (we excluded the two populations with lower plant numbers, i.e., these were not treated). Specifically, in addition to automatic watering, half of the males and half the females, respectively, received 55 mL of cytokinin and 55 ml of auxin twice a week; control plants received 55 mL of water. Following Louis and Durand (1978), we used 2 µM BAP for the cytokinin treatment and a mixture of 1 × 10□^3^ g/L IAA and 1 × 10□^2^ g/L GA□for the auxin treatment. We were not specifically interested in the effect of the hormone treatment, but rather in assessing differences in leakiness among populations. As described in the Results, the hormone treatments ultimately had no effect on flower production in either sex.

Imposition of hormone treatments necessitated separating males and females into different glasshouses. Our design thus effectively involved two independent experiments, one for each sex. We were not interested in a direct sex comparison, and we measured a different set of variables on each sex. In each glasshouse, plants were randomly distributed across seven tables in a random block design. Randomization ensured a balanced distribution of plants from each population and treatment across tables.

### Measurements of phenotypic variation

After three weeks of growth following the onset of the hormone treatments, we harvested plants table by table in each glasshouse, retaining table number and harvest time as a combined blocking unit for analysis. For most tables, all plants were harvested within two days of each other. Because of the indeterminate nature flowering in *M. annua*, our harvest represents a temporal snapshot of reproductive allocation at the time of sampling. For each plant, we recorded the number of flowers of the opposite sex (i.e., fruits on males and male flowers on females), plant height and above-ground biomass. Additionally, for a subset of five plants of each sex, population and hormone treatment (i.e., 5 x 2 x 25 x 2 for a total of 500 plants), we counted the number of female fruits on females and male flowers on males.

We measured the incidence of leakiness in a population in terms of the proportion of plants of each sex that produced at least one flower of the opposite sex. We followed Cossard and Pannell (2019) to compute the degree of leakiness for females as *d*_f_*=*λ_f_*/M*, where λ_f_ is the given female’s male flower production per biomass unit (i.e., its male RE), and *M* is the male reproductive effort of an average male from the same population. Due to phenotyping constraints, *M* for each population was calculated as the average of five untreated males selected for detailed phenotyping. The degree of leakiness for males was computed as *d*_m_*=*λ_m_*/ F*, where λ_m_ and *F* are defined analogously to λ_f_ and *M*, respectively. Dividing λ by *M* and *F* served to calibrate the degree of leakiness in one sex in relation to the typical production of flowers by plants of the opposite sex, also ensuring that values fall within the range of 0 to 1.

We calculated an index of reproductive effort, defined as the number of male flowers on a male or the number of female fruits on a female divided by its respective biomass. The reproductive effort of plants without hormone treatment was used to calculate the population average male or female reproductive effort used to compute the degree of leakiness.

### Statistical analysis

We constructed generalized linear mixed models (GLMMs) and linear models to assess the dependence of seven plant traits on distance from the assumed point of departure for the range expansion, which we refer to as the ‘expansion origin’; we set this origin as the midpoint between the two southeastern-most populations sampled (Figure 1, Table S1). The traits were: plant biomass, plant height, incidence of leakiness, degree of leakiness, and reproductive effort. We computed each site’s distance from the expansion origin (Table S1) using geodesic distance (Vincenty, 1975).

To analyse trait variation recorded in each of the (male and female) common gardens, we used GLMMs with a consistent fixed and random effect structure, differing only in response variables, error distribution, offset terms, and zero-inflation components, when applicable. We included distance from the expansion origin, hormone treatment (binary), and the interaction between distance and hormone treatment as fixed effects and population and table number (with sampling time) as random effects.

We modelled the incidence of leakiness as a binary variable and the degree of leakiness using a zero-inflated negative binomial GLMM, with counts of leaky flowers (males) or fruits (females) as the response variable, offset by the log-transformed product of plant biomass and population-average reproductive effort of the opposite sex. The zero-inflation component included distance from the expansion origin as a fixed effect, based on results from the binomial model.

Reproductive effort was modelled with a negative binomial GLMM, using counts of fruits or male flowers, offset by the logarithm of biomass. Plant height and biomass were modelled with Gaussian GLMMs; biomass was square-root-transformed.

Our model for female biomass as a function of geodesic distance failed to converge. To address this issue, we removed greenhouse table from the random effects, retaining population as the only random effect. This approach was taken because variance among tables in female biomass was effectively zero, while the variance among populations was substantial. We also compared the AIC of two models, each including only one random effect, either table or population. The model with population had a lower AIC (10,300) than the model with table number (11,200), further supporting our choice to remove table number from the final model.

All analyses were performed in R (R Core Team, 2025). All GLMMs and LMs were fitted using the ‘glmmTMB’ function from the ‘glmmTMB’ package in R (Brooks *et al*., 2017). We checked the diagnostic fit of all chosen models for significant deviations, overdispersion, and zero-inflation (when applied) using the ‘DHARMa’ package (Florian, 2016) in R, with 5,000 simulation points.

## Results

### Effect of hormone treatment on reproductive and vegetative traits

We found no effect of the hormone treatment on any of the reproductive traits studied, including leakiness ability, leakiness degree, or reproductive effort in either sex (Table S2). Hormone treatment did, however, affect plant biomass (*Z* = 9.49, *P* < 0.001 for females; *Z* = 3.57, *P* < 0.001 for males) and height (*Z* = 7.57, *P* < 0.001 for females; *Z* = 4.80, *P* < 0.001 for males), as well as significant interactions between plant size and distance from the expansion origin (Table S2). As the effect of hormones on plant size was not the focus of our study, we do not analyze plant size measurements further, except in terms of its role in the computation of reproductive effort and its variation with respect to distance from the expansion origin (Table S2).

### Greater leakiness with distance from the expansion origin

The incidence of leakiness increased in both sexes with distance from the expansion origin (females: ∼0.012 % per km from the expansion origin, *Z* = 4.29, *P* < 0.001; males: ∼0.010 % per km from the expansion origin, *Z* = 2.38, *P* < 0.05; Figure 2A, Table S2). This trend was further reflected by the zero-inflation components of models for the degree of leakiness, with the proportion of excess zeros (i.e., non-leaky individuals) declining significantly with increasing distance from the expansion origin (females: *Z* = –4.76, *P* < 0.001; males: *Z* = –3.29, *P* < 0.01; Table S2). The degree of leakiness showed no significant association with the distance from the expansion origin for either sex (Table S2).

**Figure 2.**
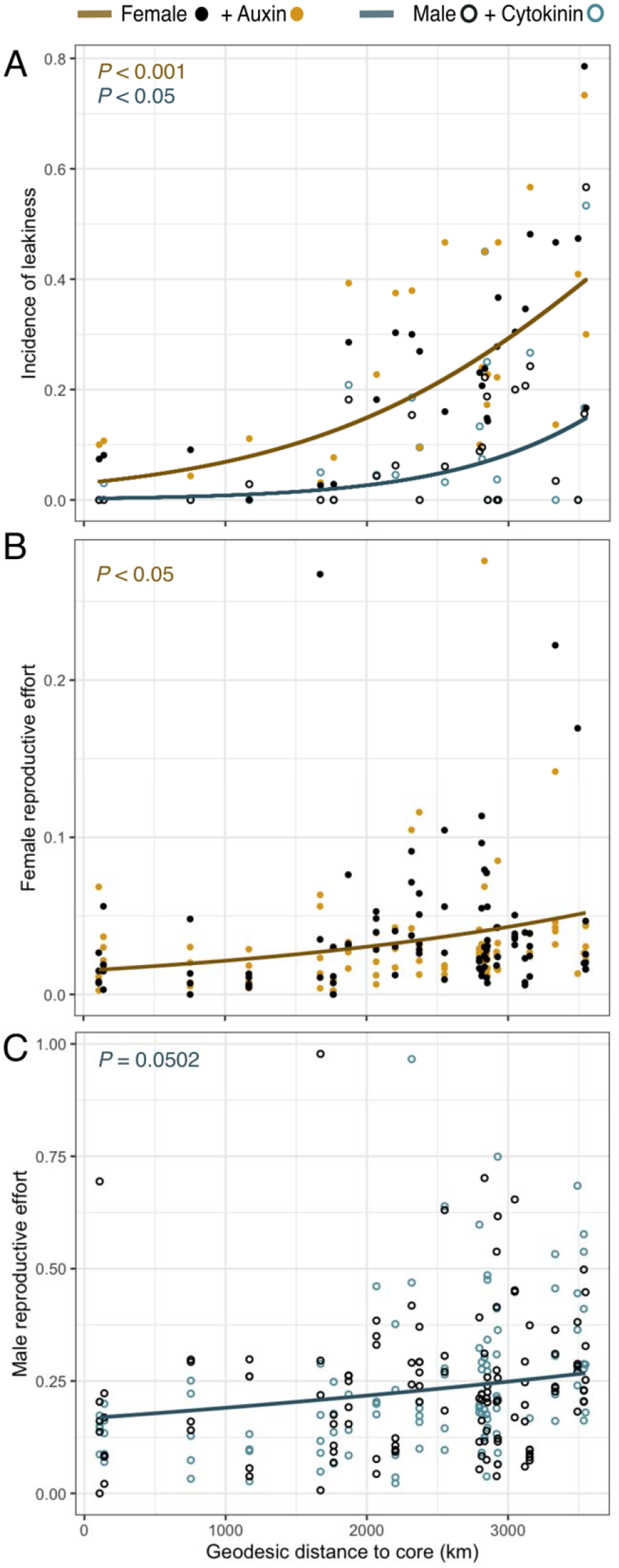
Measures of leakiness (A) and reproductive effort for females (B) and males (C) sampled as a function of distance from the expansion origin, based on traits expressed in the common-garden. Indicated are plant sex and hormone-treatment status (see inset legend). Lines show GLMM-predicted responses (see Table S2) as functions of distance from the expansion origin. Statistical significance is indicated in the upper left of each panel.

### Variation in vegetative traits with distance from the expansion origin

There was no significant association between the distance from the expansion origin and vegetative traits (biomass or height) in either sex (Table S2).

### Greater reproductive effort with distance from the expansion origin

Reproductive effort in both sexes increased with distance from the expansion origin (females: ∼8.17 x 10^-5^ flowers/g per km from the expansion origin, Z = 2.53, *P* < 0.05; males: ∼6.20 x 10^-5^ flowers/g per km from the expansion origin, *Z* = 1.96, *P* = 0.0502; Figure 2B, C, Table S2).

## Discussion

Our study reveals that both the incidence of leaky sex expression and reproductive effort in dioecious *M. annua* increase with distance from the species’ expansion origin in the eastern Mediterranean Basin and towards its geographical range margin in northern and western Europe. Both these trends are consistent with expectations for the effects of selection during a past range expansion. Of course, our results are based on correlations, not a controlled experiment, so that any causal inference needs to be cautious; in principle, other known or unknown factors could be responsible for the non-random patterns observed. For instance, the geographic gradient from southeastern to northwestern Europe coincides with substantial and partially directional changes in climate, and the observed increases in leakiness and reproductive effort along this gradient are also strongly associated with variation in climatic variables (see Supporting Information B, Table S4). Nevertheless, it is difficult to see why a temperate climate should have selected for increased leakiness. We thus discuss our results here in terms of the hypotheses that motivated our study, though the reader should bear in mind the correlational nature of our evidence.

### Increased leakiness in sex expression with range expansion

We found a significantly greater incidence of leakiness in sex expression in both males and females of *M. annua* at sites further from the expansion origin, consistent with the hypothesis that selection for reproductive assurance during repeated colonization has contributed to phenotypic evolution at the species’ range margin. Baker (1955) first suggested that the enriched capacity for uniparental reproduction on oceanic islands could be a result of selection for reproductive assurance during colonization, an idea also invoked for the evolution of a selfing ability in metapopulations (Pannell & Barrett, 1998) and range expansions (Cheptou, 2012; Pannell, 2015; Pannell *et al*., 2015). Our results here provide empirical support for this idea in the context of the selection of a capacity for self-fertilization in dioecious species via leaky sex expression.

We also expected males to show a greater increase in the degree of leakiness from the expansion origin than females, because the success of colonization by single males should be enhanced by the production of larger numbers of female flowers and seeds, whereas a single female should only produce as many male flowers as necessary to fertilize her ovules. However, our results revealed no such difference. It is possible that the advantages of a high degree of leakiness in males during colonization are countered by the disadvantages associated with producing many seeds in established populations. Because males disperse a great deal of pollen to the wind, any ovules they produce are likely to be self-fertilized, even in established populations, and self-fertilized seeds might suffer inbreeding depression (Charlesworth & Charlesworth, 1987; Pannell & Jordan, 2022).

While Baker (1967) and others (Ehlers & Bataillon, 2007; Crossman & Charlesworth, 2014; Cossard & Pannell, 2019; Cossard *et al*., 2021) have recognized that leaky sex expression in dioecious species should facilitate their colonization potential, our study now provides strong, albeit correlational, evidence that selection during repeated colonization may have enhanced this tendency in a wild plant species. An important implication is that leaky sex expression found in natural populations of dioecious species may be actively maintained by natural selection during occasional periods where mates are limited, either in low-density populations or during colonization. In other words, leaky sex expression in dioecious plants may not always be just a relic of an unfinished transition to fully separate sexes from hermaphroditism or monoecy (Lloyd, 1980) or an expression of developmental noise (Lloyd & Bawa, 1984). Indeed, dioecy in the genus *Mercurialis* is ancestral, not derived (Krähenbühl *et al*., 2002; Obbard *et al*., 2006a), and the evolution of leaky sex expression in *M. annua* can almost certainly be regarded as an evolved adaptation to its colonizing habit that coincided a transition to an annual life history (Pannell *et al*., 2008). This adaptation appears to have been enhanced not only by ongoing colonization in dynamic metapopulations (Obbard *et al*., 2006b; Eppley & Pannell, 2007b; Dorken *et al*., 2017) but also by range expansion.

### Increased reproductive effort with range expansion

We found that both male and female reproductive effort (RE) were higher toward the western range margins of *M. annua*, with female RE showing a stronger increase with distance from the expansion origin than male RE. To this extent, our results strengthen other correlative evidence for selection of traits associated with a ruderal life history during colonization in range expansions (Cwynar & MacDonald, 1987; Phillips *et al*., 2006; Darling *et al*., 2008; Monty & Mahy, 2010; Bufford & Hulme, 2021). Note that we did not find a corresponding decrease in biomass or height, as expected for a scenario of reduced competition during range expansion. Given that *M. annua* populations grow quickly after colonization (Dorken *et al*., 2017) and plants in established dense populations must be highly competitive, this result is perhaps not surprising. Trade-offs in competitive ability likely involve multiple traits (Stearns, 1983, 1989) and may produce patterns that are too weak to detect in height or biomass alone.

### Concluding remarks

Our results suggest that selection during range expansion has left a detectable phenotypic signature in *Mercurialis annua*, favouring traits likely to enhance colonization success.

Populations further from the inferred expansion origin showed a higher incidence of leaky sex expression in both sexes, consistent with the idea that reproductive assurance can be strengthened during repeated colonization, even in a dioecious species. They also showed greater reproductive effort, in line with broader life-history expectations that expansion-front populations should evolve more ruderal, colonization-oriented strategies. Although our study is correlational and cannot exclude alternative explanations, the patterns we document are nevertheless consistent with long-standing ideas derived from Baker’s Law (Baker, 1955, 1967; Stebbins, 1957; Pannell & Barrett, 1998; Cheptou, 2012; Pannell, 2015; Pannell *et al*., 2015), life-history theory (MacArthur & Wilson, 1967; Pianka, 1970; Grime, 1974; Stearns, 1983, 1989; Reznick *et al*., 2002; Travis *et al*., 2023), and models of evolution during range expansion (e.g., reviewed in Hargreaves & Eckert, 2014; Chuang & Peterson, 2016; Weiss-Lehman & Shaw, 2022). More generally, our findings suggest that the evolutionary processes associated with colonization may contribute to shaping phenotypes at species’ range margins, where mate limitation and demographic stochasticity may be pronounced. Accordingly, the evolution of traits that conferred reproductive assurance and establishment success on a species during a range expansion may then also influence not only its population and metapopulation dynamics at the range margin but also its capacity to move beyond the range margin, e.g., when a warming climate permits further range expansion.

## Acknowledgements

We thank Kai-Hsiu Chen, Shengman Lyu, Antoine Guissan, Valentin Verdon for their comments on our analysis; Leonardo Cantoro, Sebastian Cato, Theophanis Constantinidis, Stefano Doglio, Galina Gavrilenco, Thijs Heijman, Gabriele La Grasta, Ramy Maalouf, Arian Ndoni, Thomas Schorr, Gertraud Seiser, Kathy Vivian, Jean-Paul Wolff, Walter Wimmer for their help with seed sampling; Jessica Bainbridge, Kai-Hsiu Chen, Camilo Ferron Martinez, Ivan Giordano, Xia Jiang, Ehouarn Le Faou, Marjolaine Rousselle, Philipp Schenkel, as well as numerous undergraduate assistants, for their help in setting up experiments and measuring plant phenotypes; and the Swiss National Science Foundation (Grant 31003A_163384) and the University of Lausanne for funding.

