## Supporting Information for "Geographical gradients in leaky sex expression and reproductive effort in a dioecious plant are consistent with selection during range expansion"

1015 Lausanne

Switzerland

**Supporting Information**

This file comprises two sections. Section A provides additional information supporting the main analyses presented in the manuscript, namely GPS coordinates of sampling locations, sample sizes, and common-garden sex ratios, and the full results of the GLMMs reported in the main text, i.e,. the relationships between the incidence and degree of leakiness, reproductive effort, biomass, and height, and distance from the expansion origin.

Section B provides supplementary analyses examining the associations between local climate at collection sites and the traits studied in the main text, including methods and results. It also compares the relative importance of local climate and distance from the expansion origin in explaining variation in these traits.

**A. Supporting information for main analyses**

**Table S1.** The site ID and coordinates of the 26 sampling sites and the range core, the distances of each site to the core, the number of collected field mothers and repotted plants in the experiment. Note that seeds from sites 7, 11 and 13 did not germinate and are therefore not included.

| **ID** | **N (degree)** | **E (degree)** | **Geodesic distance (km)** | **Collected field mothers** | **Repotted females** | **Repotted males** |
| --- | --- | --- | --- | --- | --- | --- |
| 1 | 49.83 | 5.95 | 3047 | 30 | 22 | 21 |
| 2 | 46.52 | 6.61 | 2848 | 91 | 60 | 60 |
| 3 | 51.53 | -0.46 | 3536 | 30 | 60 | 60 |
| 4 | 36.92 | 15.10 | 1871 | 30 | 56 | 57 |
| 5 | 47.93 | 7.76 | 2833 | 30 | 39 | 40 |
| 6 | 48.20 | 16.37 | 2317 | 24 | 56 | 56 |
| 8 | 51.15 | 5.92 | 3119 | 30 | 27 | 28 |
| 9 | 33.94 | 35.62 | 141 | 30 | 60 | 60 |
| 10 | 51.53 | 0.24 | 3492 | 30 | 43 | 44 |
| 12 | 52.10 | 10.42 | 2919 | 30 | 53 | 53 |
| 14 | 45.56 | 10.20 | 2551 | 21 | 60 | 60 |
| 15 | 41.92 | 12.80 | 2202 | 14 | 48 | 48 |
| 16 | 43.76 | 15.77 | 2068 | 30 | 44 | 44 |
| 17 | 43.15 | 5.71 | 2797 | Bulk > 30 | 60 | 60 |
| 18 | 43.21 | 5.52 | 2813 | Bulk > 30 | 53 | 53 |
| 19 | 40.42 | 17.62 | 1765 | Bulk > 30 | 59 | 59 |
| 20 | 36.38 | 28.03 | 754 | 30 | 60 | 60 |
| 21 | 37.97 | 23.79 | 1169 | 30 | 60 | 60 |
| 22 | 41.32 | 19.45 | 1672 | 5 | 54 | 54 |
| 23 | 46.53 | 6.53 | 2854 | Bulk > 30 | 47 | 49 |
| 24 | 48.86 | 16.24 | 2372 | Bulk > 30 | 42 | 42 |
| 25 | 51.33 | 9.37 | 2927 | Bulk > 30 | 60 | 60 |
| 26 | 44.76 | -0.66 | 3334 | Bulk > 30 | 45 | 46 |
| 27 | 31.82 | 35.11 | 108 | Bulk > 30 | 56 | 56 |
| 28 | 51.92 | 6.08 | 3153 | Bulk > 30 | 60 | 60 |
| 29 | 50.82 | -1.09 | 3548 | Bulk > 30 | 60 | 60 |
| Core | 32.78 | 35.01 | 0 |  |  |  |

**Table S2.** Estimates for fixed effects of GLMMs. Estimates are presented with 95% confidence intervals. Significance codes : 0 *** 0.001 ** 0.01 * 0.05. In bold are estimations of fixed factors with the significance levels of less than or equal 0.05.

| Trait ~ Fixed terms + (1\| Population ID) + (1\| Table ID) | | | | | | |
| --- | --- | --- | --- | --- | --- | --- |
|  | Incidence of leakiness | | | Reproductive effort | | |
|  | Male | Female | | Male | | Female |
| Intercept | -5.99 *** (-8.55, -3.42) | -3.46 *** (-4.50, -2.43) | | -1.79 *** (-2.16, -1.42) | | -4.19 *** (-4.79, -3.58) |
| Geodesic distance | **1.19 x 10^-3^ * (0.21 x 10^-3^, 2.18 x 10^-3^)** | **8.61 x 10^-4^ *** (4.67 x 10^-4^, 12.5 x 10^-4^ )** | | **1.33 x 10^-4^ (0.00, 2.67 x 10^-4^)** | | **3.48 x 10^-4^ * (0.78 x 10^-4^, 6.17 x 10^-4^)** |
| Hormone treatment | 0.75 (-2.67, 4.16) | 0.67 (-0.55, 1.89) | | -0.354 (-0.771, 0.063) | | 2.46 (-1.52, 1.53) |
| Geodesic distance x Hormone treatment | -0.19 x 10^-3^ (-1.55 x 10^-3^, 1.16 x 10^-3^) | -1.79 x 10^-4^ (-6.78 x 10^-4^, 3.19 x 10^-4^) | | 1.45 x 10^-4^ (-0.20 x 10^-4^, 3.10 x 10^-4^) | | -6.90 x 10^-5^ (-83.4 x 10^-5^, 69.6 x 10^-5^) |
|  | Male’s biomass | Female’s biomass | | Male’s height | | Female’s height |
| Intercept | 36.4 *** (26.9, 45.9) | 39.7 *** (26.3, 53.1) | | 23.2 *** (18.5, 29.9) | | 23.6 *** (17.2, 29.9) |
| Geodesic distance | 0.80 x 10^-3^ (-2.87 x 10^-3^, 4.47 x 10^-3^) | 2.78 x 10^-3^ (-2.43 x 10^-3^, 8.00 x 10^-3^) | | 0.11 x 10^-3^ (-1.71 x 10^-3^, 1.93 x 10^-3^) | | 0.68 x 10^-3^ (-1.77 x 10^-3^, 3.13 x 10^-3^) |
| Hormone treatment | **4.30 ** (1.73, 6.87)** | **10.8 *** (7.54, 14.1)** | | **2.31 ** (0.83, 3.79)** | | **4.50 *** (2.68, 6.32)** |
| Geodesic distance x Hormone treatment | -0.93 x 10^-3^ (-1.95 x 10^-3^, 0.08 x 10^-3^) | **-1.85 x 10^-3^ ** (-3.15 x 10^-3^, -0.546 x 10^-3^)** | | -3.68 x 10^-4^(-9.53 x 10^-4^, 2.17 x 10^-4^) | | **-0.77 x 10^-3^ ***  **(-1.50 x 10^-3^, -0.05 x 10^-3^)** |
| Trait ~ Fixed terms + (1\| Population ID) + (1\| Table ID), zero inflation ~ Distance | | | | | | |
|  | Degree of leakiness in males | | Degree of leakiness in females | | | |
|  | Conditional model | Zero-inflation model | Conditional model | | Zero-inflation model | |
| Intercept | -6.85 *** (-10.7, 3.03) | 2.94 ** (1.05, 4.84) | -6.55 *** (-7.83, -5.28) | | 1.50 ** (0.55, 2.45) | |
| Geodesic distance | 6.16 x 10^-4^ (-7.08 x 10^-4^, 19.4 x 10^-4^) | **-1.22 x10^-3^ ** (-1.94 x10^-3^, -0.49 x10^-3^)** | 3.33 x 10^-4^ (-1.11 x 10^-4^, 7.78 x 10^-4^) | | **-9.55 x 10^-4^ ***(-13.5 x 10^-4^, -5.61 x 10^-4^)** | |
| Hormone treatment | 1.36 (-1.50, 4.22) | - | 0.29 (-0.97, 1.55) | | - | |
| Geodesic distance x Hormone treatment | -5.27 x 10^-4^ (-14.9 x 10^-4^ , 4.39 x 10^-4^) | - | -1.43 x 10^-4^ (-5.90 x 10^-4^, 3.05 x 10^-4^) | | - | |

**B. Climatological analyses**

Methods

Given that plant traits might have been influenced not only by the history of range expansion but also by environmental factors and local adaptation especially to climate after colonisation, we asked whether the significant patterns we observed across the species range (in terms of the incidence of leakiness and reproductive effort; see Results) remain significant when climatological factors are included in the models. To do this, we first analyzed the climatological data for the populations and then rebuilt models for these focal traits, incorporating climate factors as fixed effects while excluding all non-significant terms from the previous models.

To characterize the overall local climates of our sampling sites, we extracted climatological data from 1981 to 2010 using the CHELSA global climate dataset (version 2.1; Karger *et al.*, 2017). After cleaning the dataset and removing missing values, we retained 65 climatological variables (Table S6), across 26 populations. We then used principal component analysis (PCA) on the scaled values of these variables to reduce the data dimensions to two principal components (PCs), which together explained approximately 70% of the total climatic variance (PC1: 54.7%; PC2: 14.2%). We tested for pairwise correlations among the two distance metrics and the two climate PCs using Pearson correlation tests.

We next built generalized linear mixed models (GLMMs) for the incidence of leakiness and reproductive effort using geodesic distance from the species core and the two climate PCs as fixed effects. These models were selected based on previous analyses that showed significant associations between these traits and distance. All hormone-related terms, previously found to be non-significant, were excluded from these models. Random effects, data distribution families, and offset terms were retained, consistent with the previous models.

To evaluate the relative contributions of distance from the species core and local climate on trait variation, and to address multicollinearity arising from the inclusion of two distance metrics and two climate PCs in the same model, we applied variation partitioning we conducted commonality analysis (Ray-Mukherjee *et al.*, 2014), i.e., variation partitioning (Goldstein *et al.*, 2002), for all the models of these four traits, using geodesic distance and two climate PCs as fixed effects. We compared the marginal R^2^ (Nakagawa *et al.*, 2017), the proportion of variance in the response variable being explained by the fixed factors, for full models with models with subsets of fixed effects, quantifying variance explained by each factor. Additionally, we also performed analogous analyses using land distance instead of geodesic distance, and constructed GLMMs using climate principal components (PCs) in place of distance from the core, and variation partitioning was performed using both distance metrics and climate PCs.

Correlation tests were performed using the ‘cor’ function from the ‘stats’ package (Bolar, 2019). Variation partitioning followed the method in Legendre (2008), modified for the higher number of fixed factor in our models. Marginal R^2^ of the models were extracted using the ‘performance’ package in R (Lüdecke *et al.*, 2021).

Results

The PC1 scores for local climate were moderately to highly correlated with geodesic distance (*r* = 0.82), reducing the predictive power of each factor in models that included both geodesic distance and climate PC1, and complicating interpretation when only a subset of the correlated terms was included. When both principal components (PCs) of local climate were added to the models, the associations between the incidence of leakiness and geodesic distance remained positive but fell short of statistical significance (males: *Z* = 1.91, *P* = 0.057; females: *Z* = 1.85, *P* = 0.064; Table S3). Leakiness showed no significant association with either of the climate PCs (*P* > 0.2 for both; Table S3). Geodesic distance showed no significant association with reproductive effort in either sex (Table S3).

Our analysis of variation partitioning revealed that geodesic distance was the dominant factor explaining variance in the incidence of leakiness in (Figure S1). By contrast, most of the variance in the incidence of leakiness in females and reproductive effort in both sexes was explained by the combined effects of distance and climate. Across all four focal traits, distance metric consistently explained a larger proportion of marginal variance than climate PCs (Figure S1).

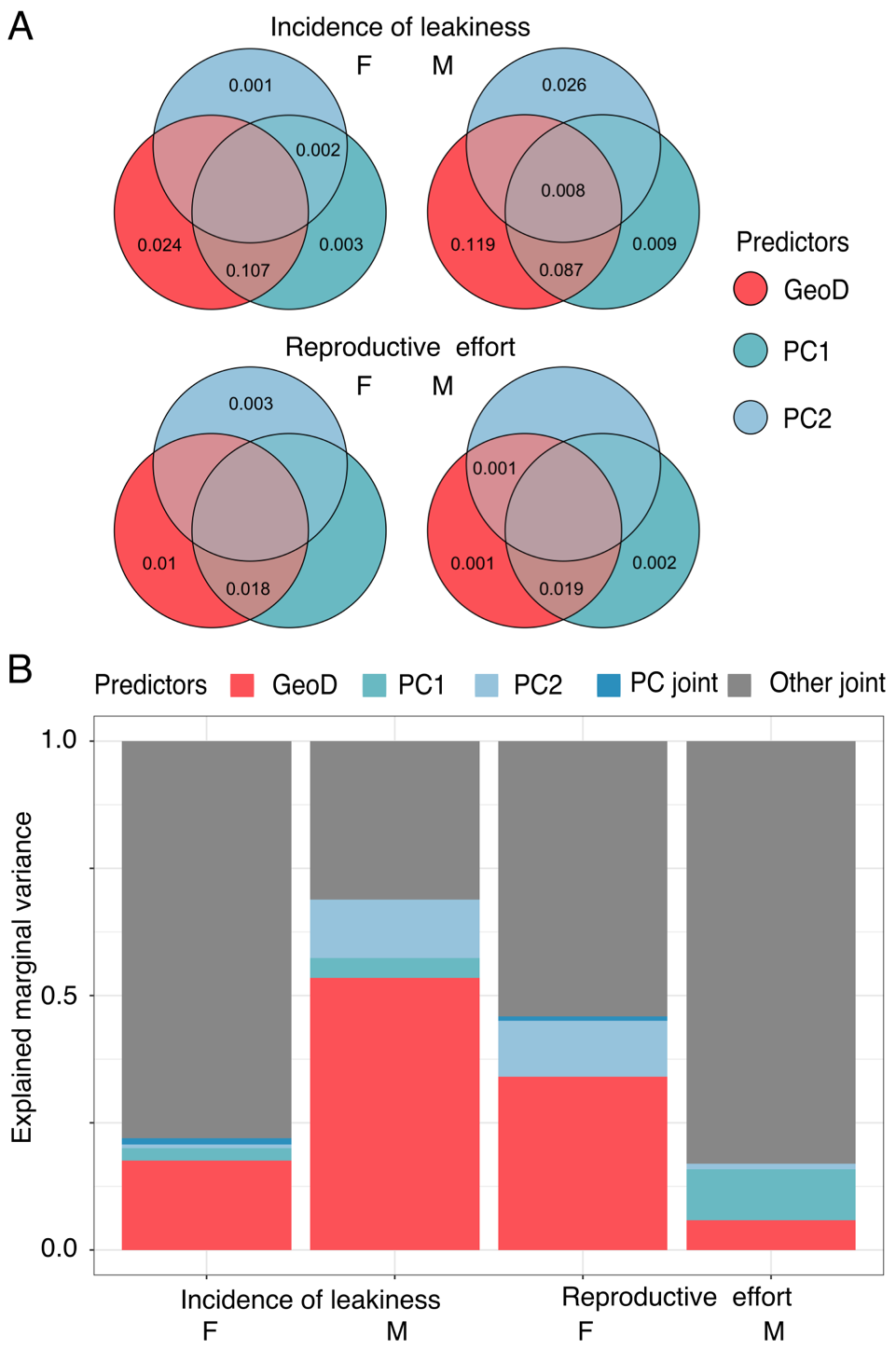

**Figure S1.** Variation partitioning of GLMMs for each trait and sex that showed significant associations with geodesic distance from the range’s core. (A) Venn diagrams showing the proportion of total variance in each response trait explained by geodesic distance (red), two climate principal components (blue shades), and by their joint effects. Empty fractions represent non-positive contributions to explained variance. (B) Stacked bar plots showing marginal variance, i.e., the proportion of variance in each response trait explained by fixed effects, partitioned into components attributed to geodesic distance (red), climate (blue shades; including PC1, PC2, and their joint effect), and the shared contribution of distance and climate (grey).

**Table S3.** Estimates for fixed effects of GLMMs and LM using geodesic distance and PCs of climate as fixed factors. Estimates are presented with 95% confidence intervals. Significance codes : 0 *** 0.001 ** 0.01 * 0.05. In bold are estimations of fixed factors with the significance levels of less than or equal 0.05.

| Trait ~ Fixed terms + (1\| Population ID) + (1\| Table ID) | | | | |
| --- | --- | --- | --- | --- |
|  | Incidence of leakiness | | Reproductive effort | |
|  | Male | Female | Male | Female |
| Intercept | -6.36 *** (-10.1, -2.63) | -2.60 *** (-4.06, -1.13) | -1.71 *** (-2.23, -1.13) | -4.27 ** (-5.83, -2.71) |
| Geodesic distance | 1.41 x 10^-3^ (-0.039 x 10^-3^, 2.86 x 10^-3^) | 5.47 x 10^-4^ (-0.325 x 10^-4^, 11.3 x 10^-4^) | 9.88 x 10^-5^ (-10.5 x 10^-5^, 30.3 x 10^-5^) | 3.48 x 10^-4^ (-3.00 x 10^-4^, 9.95 x 10^-4^) |
| PC1 | 0.046 (-0.150, 0.243) | -4.61 x 10^-2^ (-13.1 x 10^-2^, 3.91 x 10^-2^) | -1.98 x 10^-2^ (-5.25 x 10^-2^, 1.29 x 10^-2^) | 0.25 x 10^-2^ (-8.85 x 10^-2^, 9.35 x 10^-2^) |
| PC2 | 0.108 (-0.124, 0.340) | 1.77 x 10^-2^ (-8.25 x 10^-2^, 11.8 x 10^-2^) | -6.95 x 10^-3^ (-45.3 x 10^-3^, 31.4 x 10^-3^) | 0.037 (-0.036, 0.110) |

**Table S4.** Estimates for fixed effects of GLMMs and LMs using PC1 of climate. Estimates are presented with 95% confidence intervals. Significance codes : 0 *** 0.001 ** 0.01 * 0.05. In bold are estimations of fixed factors with the significance levels of less than or equal 0.05.

| Trait ~ Fixed terms + (1\| Population ID) + (1\| Table ID) | | | | | |
| --- | --- | --- | --- | --- | --- |
|  | Incidence of leakiness | | | Reproductive effort | |
|  | Male | | Female | Male | Female |
| Intercept | -3.04 *** (-3.75, -2.33) | | -1.38 *** (-1.75, -1.01) | -1.47 *** (-1.67, -1.28) | -3.37 *** (-3.65, -3.08) |
| PC1 | **-0.120 * (-0.239 , -0.001)** | | **-0.119 *** (-0.175, -0.063)** | **-0.029 * (-0.052, -0.007)** | **-0.041 * (-0.076, -0.006)** |
| Hormone treatment | 0.171 (-0.262, 0.604) | | 0.216 (-0.064, 0.496) | 0.003 (-0.154, 0.161) | -0.146 (-0.362, 0.069) |
| PC1 x Hormone treatment | -0.014 (-0.098, 0.070) | | 0.005 (-0.045, 0.057) | -0.009 (-0.037, 0.020) | -0.006 (-0.043, 0.031) |
|  | Male’s biomass | | Female’s biomass | Male’s height | Female’s height |
| Intercept | 38.3 *** (34.8, 41.8) | | 45.8 *** (40.6, 51.0) | 23.4 *** (21.7, 25.2) | 25.2 *** (22.8, 27.5) |
| PC1 | 0.134 (-0.461, 0.729) | | -0.109 (-0.97, 0.749) | 0.071 (-0.223, 0.366) | -0.026 (-0.424, 0.372) |
| Hormone treatment | **2.10 *** (1.07, 3.12)** | | **6.48 *** (5.21, 7.76)** | **1.47 *** (0.878, 2.07)** | **2.73 *** (2.02, 3.44)** |
| PC1 x Hormone treatment | -0.045 (-0.126, 0.217) | | 0.026 (-0.191, 0.243) | -0.041 (-0.140, 0.058) | -0.009 (-0.129, 0.112) |
| Trait ~ Fixed terms + (1\| Population ID) + (1\| Table ID), zero inflation ~ PC1 | | | | | |
|  | Male’s degree of leakiness | | | Female’s degree of leakiness | |
|  | Conditional model | Zero-inflation model | | Conditional model | Zero-inflation model |
| Intercept | -5.58 *** (-6.60, -4.57) | -1.15 (-3.14, 0.846) | | -5.76 *** (-6.22, -5.30) | -0.995 (-2.01, 0.023) |
| PC1 | -0.054 (-0.229 , 0.121) | 0.210 (-0.004, 0.424) | | -0.036 (-0.106, 0.035) | **0.194 *** (0.093, 0.296)** |
| Hormone treatment | -0.182 (-0.794, 0.432) | - | | -0.046 (-0.398, 0.306) | - |
| PC1 x Hormone treatment | -0.062 (-0.186 , 0.061) | - | | 0.030 (-0.035, 0.095) | - |

**Table S5.** Estimates for fixed effects of GLMMs and LMs using PC2. Estimates are presented with 95% confidence intervals. Significance codes : 0 *** 0.001 ** 0.01 * 0.05. In bold are estimations of fixed factors with the significance levels of less than or equal 0.05.

| Trait ~ Fixed terms + (1\| Population ID) + (1\| Table ID) | | | | | | | |  |
| --- | --- | --- | --- | --- | --- | --- | --- | --- |
|  | Incidence of leakiness | | | Reproductive effort | | | |  |
|  | Male | Female | | Male | | Female | |  |
| Intercept | -3.08 *** (-3.85, -2.30) | -1.38 *** (-1.82, -0.932) | | -1.47 *** (-1.69, -1.26) | | -3.37 *** (-3.67, -3.08) | |  |
| PC2 | -0.016 (-0.261, 0.229) | -0.068 (-0.204, 0.068) | | -0.002 (-0.055, 0.050) | | 0.004 (-0.069, 0.078) | |  |
| Hormone treatment | 0.192 (-0.215, 0.598) | 0.196 (-0.080, 0.471) | | -0.011 (-0.168, 0.147) | | -0.170 (-0.386, 0.046) | |  |
| PC2 x Hormone treatment | 0.043 (-0.092, 0.177) | 0.085 (-0.006, 0.176) | | -0.024 (-0.078, 0.029) | | 0.026 (-0.044, 0.096) | |  |
|  | Male’s biomass | Female’s biomass | | Male’s height | | Female’s height | |  |
| Intercept | 38.3 *** (34.7, 41.8) | 45.8 *** (40.7, 51.0) | | 23.4 *** (21.8, 25.1) | | 25.2 *** (22.9, 27.5) | |  |
| PC2 | 0.307 (-0.855, 1.47) | 0.238 (-1.42, 1.90) | | 0.465 (-0.086, 0.102) | | 0.363 (-0.389, 1.11) | |  |
| Hormone treatment | **2.13 *** (1.11, 3.15)** | **6.37 *** (5.12, 7.62)** | | **1.45 *** (0.858, 2.04)** | | **2.67 *** (1.97, 3.37)** | | - |
| PC2 x Hormone treatment | 0.155 (-0.169, 0.479) | **1.18 *** (0.778, 1.60)** | | 0.014 (-0.174, 0.202) | | **0.470 *** (-0.241, 0.699)** | | - |
| Trait ~ Fixed terms + (1\| Population ID) + (1\| Table ID), zero inflation ~ PC2 | | | | | | | | |
|  | Male’s degree of leakiness | | | | Female’s degree of leakiness | | | |
|  | Conditional model  (Non-converged) | | Zero-inflation model  (Non-converged) | | Conditional model | | Zero-inflation model | |
| Intercept | -6.10 | | -19.35 | | -6.06 *** (-6.54, -5.59) | | -2.58 ** (-4.39, -0.759) | |
| PC2 | -0.097 | | -0.666 | | 0.019 (-0.130, 0.167) | | **0.415 * (0.089, 0.741)** | |
| Hormone treatment | -0.084 | | - | | -0.105 (-0.429, 0.219) | | - | |
| PC2 x Hormone treatment | -0.145 | | - | | 0.077 (-0.032, 0.187) | | - | |

**Table S6.** Climatological variables used for the analysis of local climates at the 26 sites sampled.

| Variable names - CHELSA database | Description |
| --- | --- |
| bio1 | Mean annual temperature calculated as the average of mean monthly temperatures over the year |
| bio2 | Mean diurnal temperature range computed as the average of monthly |
| bio3 | Isothermality; compares day–night variability to annual temperature range |
| bio4 | Temperature seasonality given by the standard deviation of mean monthly temperatures |
| bio5 | Highest monthly mean of daily maximum temperatures across the year; indicates peak thermal conditions |
| bio6 | Lowest monthly mean of daily minimum temperatures across the year; characterizes winter cold intensity |
| bio7 | Annual temperature range; measures amplitude between warmest and coldest months |
| bio8 | Average monthly mean temperature over the wettest 3-month period of the year |
| bio9 | Average monthly mean temperature over the driest 3-month period of the year |
| bio10 | Average monthly mean temperature over the warmest 3-month period of the year |
| bio11 | Average monthly mean temperature over the coldest 3-month period of the year |
| bio12 | Sum of monthly precipitation totals across the year |
| bio13 | Maximum monthly precipitation total |
| bio14 | Minimum monthly precipitation total |
| bio15 | Coefficient of variation of monthly precipitation totals |
| bio16 | Average monthly precipitation during the wettest 3-month period of the year |
| bio17 | Average monthly precipitation during the driest 3-month period of the year |
| bio18 | Average monthly precipitation during the warmest 3-month period of the year |
| bio19 | Average monthly precipitation during the coldest 3-month period of the year |
| clt_max | Highest total cloud cover percentage |
| clt_mean | Average total cloud cover percentage |
| clt_min | Lowest total cloud cover percentage |
| clt_range | The difference between highest and lowest total cloud cover percentage |
| cmi_max | Highest monthly climate moisture index |
| cmi_mean | Average monthly climate moisture index |
| cmi_min | Lowest monthly climate moisture index |
| cmi_range | The difference between highest and lowest monthly climate moisture index |
| gdd0 | Sum of daily mean temperatures above 0 °C accumulated over the year |
| gdd5 | Sum of daily mean temperatures above 5 °C accumulated over the year |
| gdd10 | Sum of daily mean temperatures above 10 °C accumulated over the year |
| gsl | Number of days between the first and last occurrence of growing season conditions |
| gsp | Total precipitation accumulated during the growing season period |
| gst | Average daily mean temperature over all growing season days |
| hurs_max | Highest near-surface relative humidity |
| hurs_mean | Average  near-surface relative humidity |
| hurs_min | Lowest near-surface relative humidity |
| hurs_range | The difference between highest and lowest near-surface relative humidity |
| kg0 | Köppen–Geiger climate classification |
| kg1 | Köppen–Geiger climate classification without As/Aw differenciation |
| kg2 | Köppen–Geiger climate classification after Peel et al. 2007 |
| kg3 | Climate classification after Wissmann 1939 |
| kg4 | Climate classification after Thornthwaite 1931 |
| kg5 | Climate classification after Troll-Pfaffen |
| ngd0 | Total number of days in a year with mean daily temperature above 0 °C |
| ngd5 | Total number of days in a year with mean daily temperature above 5 °C |
| ngd10 | Total number of days in a year with mean daily temperature above 10 °C |
| npp | Production of carbon' means the production of biomass expressed as the mass of carbon which it contains |
| pet_penman_max | Highest monthly potential evapotranspiration |
| pet_penman_mean | Average monthly potential evapotranspiration |
| pet_penman_min | Lowest monthly potential evapotranspiration |
| pet_penman_range | The difference between highest and lowest monthly potential evapotranspiration |
| rsds_max | Highest surface downwelling shortwave flux in air |
| rsds_mean | Average surface downwelling shortwave flux in air |
| rsds_min | Lowest surface downwelling shortwave flux in air |
| rsds_range | The difference between highest and lowest surface downwelling shortwave flux in air |
| scd | Number of days per year with snow cover present at the surface |
| sfcWind_max | Highest near-surface wind speed |
| sfcWind_mean | Average near-surface wind speed |
| sfcWind_min | Lowest near-surface wind speed |
| sfcWind_range | The difference between highest and lowest near-surface wind speed |
| swb | Total water equivalent of snowpack accumulated over the year |
| vpd_max | Highest vapor pressure deficit |
| vpd_mean | Average vapor pressure deficit |
| vpd_min | Lowest vapor pressure deficit |
| vpd_range | The difference between highest and lowest vapor pressure deficit |
